# Integrative analysis of transcriptome and metabolism reveals functional roles of redox homeostasis in low light and salt combined stress in *Leymus chinensis*

**DOI:** 10.1101/2024.12.04.626903

**Authors:** Jikai Li, Suyang Fang, Hailing Zhang, Zubair Iqbal, Chen Shang, Weibo Han, Kai Huang, Xiangshen Meng, Muyuan Dai, Zhiheng Lu, Bingnan Guo, Mingnan Qu

## Abstract

Soil salinity (Salt) is one of the major limiting factors of *Leymus chinensis* (named sheepgrass) performance and alters their primary metabolic process and gene expression. Salt stress could accelerate inhibitive effects concomitant with low light (LL-Salt). However, little is known about physiological and molecular mechanisms under such LL-Salt in sheepgrass. This study aims to uncover the key reprogrammed metabolic pathways induced by LL-Salt through an integrated analysis of transcriptome and metabolism. Plant seedlings were exposed to six combinations of light intensity (moderate light, ML, and LL) and NaCl concentrations (0, 50, and 200mM) for 20 days. Results suggest that the growth of sheepgrass seedlings was dramatically inhibited with severe lodging phenotype, especially when exposed to LL-Salt combined conditions. Activities of an antioxidant enzyme, catalase, were significantly increased in LL but significantly decreased in salt stress, leading to a reverse pattern for H2O2 in LL and salt stress. Transcriptome analysis reveals 4921 downregulated differentially expressed genes (DEGs) with carbon metabolism pathways were significantly enriched compared to light response without salt (ML_0 vs LL_0). There are 194 overlapped downregulated DEGs induced by two salt treatments (ML_50 vs ML_0 and ML_200 vs ML_0), where the carbon metabolism pathway was also significantly enriched. In terms of interactive effects of LL-Salt treatments, we found that there are 16 overlapped DEGs, including a phytochrome-interacting transcription factor 4 (*PIF4*) with upregulation in LL treatment while downregulation in salt treatment, which was validated by qPCR. Metabolism analysis confirmed that in carbon metabolism, the contents of seven differentially abundant metabolites (DAMs), including sucrose, were all downregulated in both LL and salt treatment. In contrast, serine and glycolate in the photorespiration pathway were downregulated in LL while upregulated in salt treatment. A photorespiration regulatory gene, glycolate oxidase expression, was confirmed by qPCR. Collectively, we found that serval antioxidant redox pathways, including photorespiration, GSG/GSSH redox, and ABA signaling, participated in response to LL and salt combined events and highlighted the roles of cellular redox homeostasis in LL-Salt response in sheepgrass.

## Introduction

Plants are constantly exposed to shade conditions due to many natural factors, such as high planting density, cloudy weather, smog, and self-shading. The agricultural importance of long-term shade is due to its deteriorative effect on crop yield [1]. *Leymus chinensis*, commonly known as sheepgrass or Chinese ryegrass, is a perennial species of the Gramineae family [2]. Due to its high productivity and high protein content, this species is becoming major gramineous forage in Northern China and the Mongolian plateau, and it is a candidate grass extensively applied for the establishment or renewal of artificial grassland [3]. However, due to growth environments, it was constantly exposed to highly salt soil conditions. Following the increasing demands of growing sheepgrass, high-density planting directly resulted in low light (LL) conditions. Therefore, LL becomes a non-negligible factor in reducing production, especially under LL concomitant with high salt conditions (LL-Salt). However, studies on physiological and molecular regulation in response to such LL-Salt combined events were less reported, but the responsive mechanism in their separate pathway is well known.

In general, salt stress inhibits plant growth in two ways: 1) increased osmotic stress (early response), which makes it difficult for the plant to absorb water; 2) enhanced ion toxicity (late response), which is due to the effect of sodium ions on cellular functions and can affect nutrient uptake, enzyme activity, photosynthesis, and metabolism [4] Therefore, plants have adapted through various mechanisms to counteract the adverse effects of salt stress, such as inhibiting ROS production, enhancing antioxidant enzyme activities, accelerating carbon assimilation metabolism, controlling photorespiration, and mediating GSG/GSSH redox system and plant hormones [5]. Many physiological and molecular mechanisms influence these two responses, including osmotic, ion, and tissue tolerance [6].

In terms of light response, there are many ways in which plants adapt to ambient light at different levels, from the molecular to the morphological [7–9]. Continuous changes in light conditions or extremely low or high light intensity, along with other stressors such as cold, heat, and drought, are the most challenging for plants, which respond dynamically to these conditions [10]. Low light events lead to energy deficiency, and plants adapt to the resulting conditions by expanding their life forms and rearranging their leaves. High light intensities damage plants by overstimulating their chloroplasts and generating chemical reactions of free radicals. These radicals readily react with oxygen to produce reactive oxygen species (ROS) that destroy the structure of membranes, proteins, and other membrane components such as chlorophyll. However, how plants’ morphological, physiological, and molecular regulator characteristics respond to low light intensities and salinity combined stress conditions remains unclear.

In this study, we combined a comparative transcriptome and non-targeted metabolism to uncover the key molecular regulators or metabolic pathways underlying the response of sheepgrass to an LL-Salt combined event. Six combinations of light (moderate light and low light) and salt stress (0mM, 50mM, and 200mM NaCl) were performed in sheepgrass seedlings. We systematically investigated the morphological and physiological traits in sheepgrass within 20 days of LL-Salt. Based on transcriptome analysis, GO and KEGG analysis suggest that many overlapped biological pathways were significantly enriched in light and salt response, but some have different regulatory patterns. In addition, metabolism analysis suggests a robust list of several overlapped metabolites that could be involved in the LL and salt stress response. Finally, we summarized the key metabolic pathways that co-respond to LL and salt stress and highlighted the important role of redox homeostasis in response to both LL and salt stress in sheepgrass. This study may provide insight into how plants mediate cellular redox homeostasis to maintain growth development in such LL-Salt combined events in sheepgrass, which helps to guide precise molecular assistant selection in sheepgrass.

## Materials and methods

### Plant materials and LL-Salt treatments

Plump seeds of L. *chinensis* were sterilized with 5% sodium hypochlorite for 5 min and washed 4 times in sterile distilled water for 12 h at room temperature and then transferred to wet filter paper to germinate at room temperature (22–25 °C) for 24 h. The uniformly germinated seeds were selected to grow in plastic pots containing Hoagland solution that was changed every two days. After 3 weeks, the plants were treated with NaCl in three concentrations, 0 mM, 50 mM, and 200 mM, for 0, 2, 5, 10, and 20 days. Simultaneously, sheepgrass seedlings were exposed to two light treatments, i.e., moderate light (ML, 500 μmol m^−2^s^−1^ PPFD) and low light (LL, 50 μmol m^−2^s^−1^ PPFD). Therefore, six-light and salt treatment combinations exist, including ML_0, ML_50, ML_200, LL_0, LL_50, and LL_200. After that, partially sampled leaves were used to measure the physiological parameters immediately, and the remaining leaves were kept frozen at – 80 °C for later use.

### Physiological parameters measurement

Soluble sugars and sucrose were measured as previously reported [11]. Proline concentrations were estimated using the ninhydrin reaction method [12]. Catalase (CAT), malondialdehyde (MDA), and peroxidase (POD) enzyme activities were measured by using the kits (Cat. nos. XG6, EY2, and FY3) supplied by Suzhou Keming Science and Technology Co., Ltd. (China). Superoxide dismutase (SOD) enzyme activity was determined by the kit from the Nanjing Jiancheng Bioengineering Institute of Jiangsu Province, China (Cat. no. A001-3). Three biological replicates were used to minimize experimental error. The student’s t-test determined the statistical significance of the differences using the R package (2.3.2) software.

### Maximal Quantum Yield

To estimate the photosynthetic capacity of sheepgrass induced by LL-Salt, we measured maximal quantum yield (*F*_v_/*F*_m_) values. A Multi-Function Plant Efficiency Analyzer chlorophyll fluorometer (Hansatech) was used to measure *F*_v_/*F*_m_ following [13]. F_m_ represents the maximum chlorophyll fluorescence; F_o_ is the minimum chlorophyll fluorescence, and *F*_v_ = *F*_m_×2×*F*_0_ [14].

### Transcriptome analysis

Leaf samples were collected from sheepgrass exposed to either LL or two salt treatments. Total RNA extraction was conducted using TRIzol reagent (Invitrogen, Carlsbad, CA). RNA degradation and contamination were monitored using 1% agarose gel electrophoresis, while purity was assessed using the Nano-Photometer spectrophotometer (IMPLEN, CA, USA). The RNA Nano 6000 Assay Kit on the Agilent Bioanalyzer 2100 system (Agilent Technologies, CA, USA) was employed to evaluate RNA integrity. Subsequently, 1.5 μg of RNA per sample was used for RNA sample preparations. The NEB Next Ultra RNA Library Prep Kit for Illumina (NEB, USA) was utilized to generate sequencing libraries [15]. Following cluster generation, the prepared libraries were sequenced on an Illumina Hiseq 4000 platform, producing 150 bp paired-end reads.

The quality assessment of RNA-seq data was performed using FastQC software. Following generating the genome index, clean RNA-seq reads were aligned using STAR [16], with the ‘—quantMode GeneCounts’ option utilized to quantify the number of reads per gene. Subsequently, gene and isoform quantification was carried out using cufflinks v.2.2.1. Differentially expressed genes (DEGs) analysis among six combinations of light and salt in sheepgrass leaves was performed using the fragment per kilobase of transcript per million mapped reads (FPKM) method to assess transcript abundance. DEGs were identified employing the R package ‘DESeq2’ [17] with read counts obtained from STAR [16] considered in the analysis. Only genes exhibiting an adjusted *P*-value <0.05 were considered as DEGs. To minimize transcriptional noise, each isoform/gene was included for analysis if its FPKM values was > 0.01, based on a threshold established through gene coverage saturation analysis [18].

### qRT-PCR analysis

To further validate the expression pattern of genes induced by LL-Salt in sheepgrass, we conducted qRT-PCR analysis. The top fully expanded leaves from each plant, exposed to six combinations (ML-0, ML-50, ML-200, LL-0, LL-50, LL-200), were collected for qRT-PCR analysis. Total mRNA extraction was performed using TRIzol Reagent (Invitrogen), with subsequent removal of genomic DNA by DNase I treatment (Takara). The extracted RNA was reverse transcribed into cDNA using the SuperScript VILO cDNA Synthesis Kit (Invitrogen Life Technologies). We employed SYBR Green PCR Master Mix (Applied Biosystems, USA, 4309155) for qRT-PCR analysis on a Real-Time PCR System (ABI StepOnePlus, Applied Biosystems, USA). Primers for qRT-PCR were designed using Primer Prime Plus 5 Software v. 3.0 (Applied Biosystems, USA), and the *Actin1* gene was used as an internal reference. The relative expression of the gene against *Actin1* was quantified using the 2^−ΔΔCT^ method (ΔΔCT = CT, gene of interest^−CT^) [19]. This analysis was performed using three biological replicates. The primers are listed in Table S1.

### Metabolism analysis

Non-targeted metabolic profiling was conducted on the leaves of sheepgrass subjected to the combined LL-Salt treatment using LC-MS/MS (Q Exactive, Thermo Scientific). Approx. 20 mg leaf samples from sheepgrass were collected in a 2 ml Eppendorf tube containing pre-cooled metal beads and immediately stored in liquid nitrogen [20]. The samples were first extracted with a ball mill at 30 *hz* for 5 min, and then the extracted powder was dissolved in 1.5 ml methanol/chloroform mixture and incubated at -20 ^°C^ for 5 h. The mixture was centrifuged at 2,000 *g* and 4 ^°C^ for 10 min and eventually filtered with 0.43 μm organic phase medium (GE Healthcare, 6789-0404). The column was a 100×2 mm Phenomenex Luna 3 μm NH_2_ (Catalog No.: 00D-4377-B0, 0.314 mL volume). The injection volume was 20 μL, and the column temperature was 20 ^°C^ with 0.4 mL/min flow speed. Eluent A: 10 mM Ammonium acetate and 5% (v/v) acetonitrile solution, adjusted to pH 9.5 with ammonia water. Eluent B: acetonitrile. The following gradient was used for elution: 0–1 min, 15% A; 1–8 min, 15–70% A; 8–20 min, 70–95% A; 20–22 min, 95% A; 22–25 min, 15% A. The metabolomic analysis employed metabolon software (Durham, NC, USA). For the identification of metabolic compounds in each sample, the mass spectra from the NIST02 and the Golm metabolome database entries were considered (http://csbdb.mpimp-golm.mpg.de/csbdb/gmd/gmd.html).

## Results

In this study, we investigated the performance of sheepgrass seedlings exposed to six combinations of light (ML and LL) and salt (0, 50 and 200mM NaCl) treatments, i.e., ML-0, ML-50, ML-200, LL-0, LL-50, LL-200 for 20 days. Results suggest that the growth of sheepgrass seedings was dramatically inhibited either under LL or high salt treatment (200 mM NaCl). In particular, the plants exhibit a more severe lodging phenotype when exposed to LL-Salt combined conditions than under other conditions (Figure 1A). Inconsistent with this, *F*_v_/*F*_m_, representing an indicator of photosynthetic capacity, shows at least a 15% reduction (*P*<0.05) under LL across three NaCl concentrations compared to that under ML condition (Figure 1B). Salt could induce *F*_v_/*F*_m_ reduction even under 50mM NaCl, suggesting the NaCl treatments for 20 days are sufficient to inhibit sheepgrass growth. In addition, total sugar content, H_2_O_2,_ and proline contents were all increased under salt stress irrespective of light treatment, except for H_2_O_2_ under ML-0 condition (Figure 1C-E). This evidence suggests two different responses for the antioxidant reaction between ML and LL, although proline contents consistently increased following increased NaCl.

**Figure 1.**
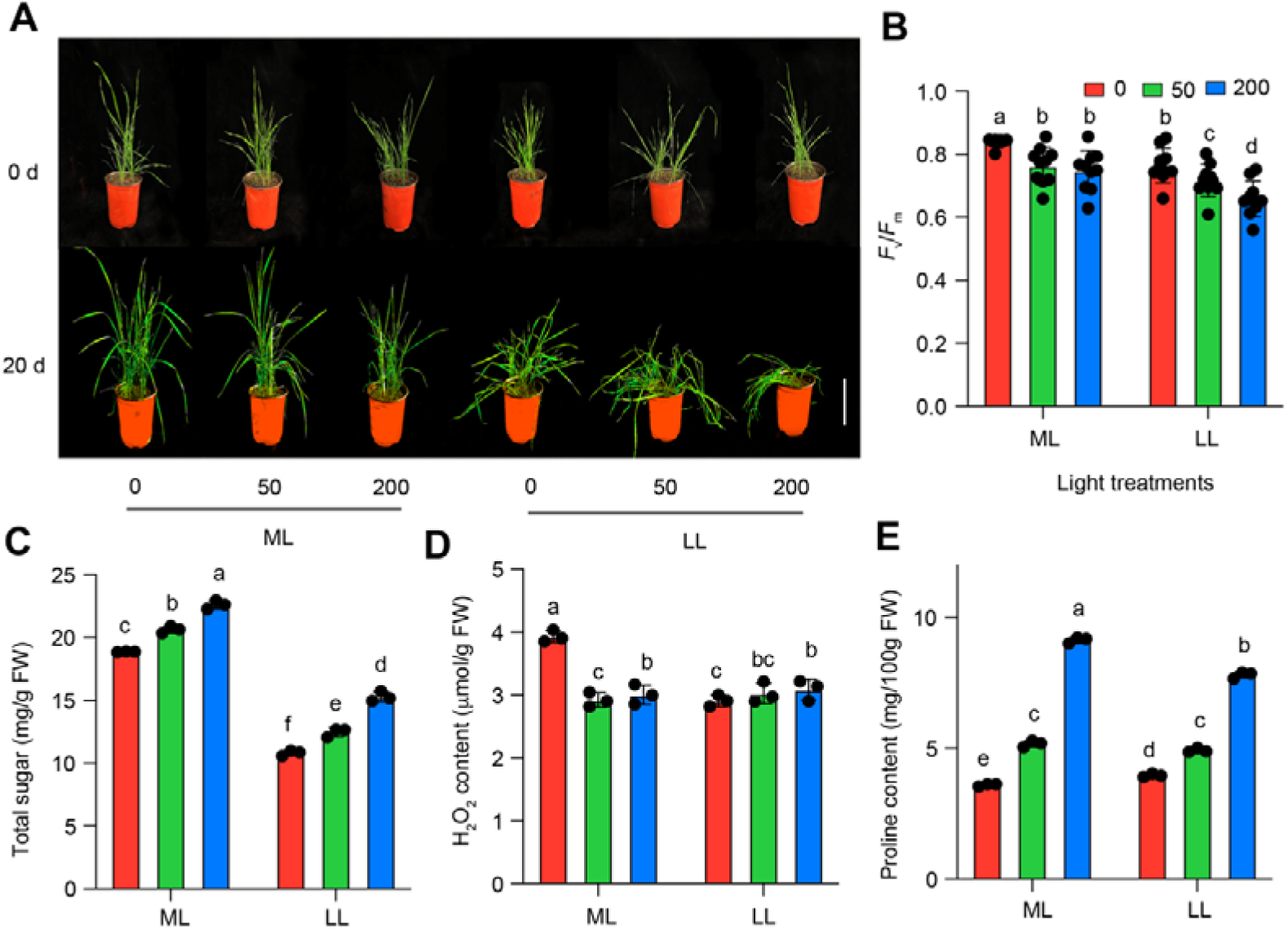
Performance of sheepgrass seedlings exposed to LL-Salt condition. **A**, Images of sheepgrass plants exposed to LL-Salt condition for 0 and 20 days. The vertical bar represents 5cm. **B-E**, *F*_v_/*F*_m_, total sugar content, H_2_O_2_ content, and proline content. Each bar data represents the mean of replicates (*n*=10 for panel **B** and *n*=4 for panels **C-E**). Different letters represent significant differences based on one-way *ANOVA* followed by Tukey’s HSD tests (*P*<0.01).

To further examine the differences in antioxidant reaction between two light conditions, we measured the enzyme actives of SOD, CAT, and MDA. Results suggest that Activities of CAT and POD were decreased following increased NaCl treatment, while they are opposite for MDA and SOD activity (Figure S1A-D). In terms of light treatment with a comparison of ML-0 vs LL-0, there were different changes for the four antioxidant enzymes, with an increase in CAT and SOD, a decrease in POD, and no significant difference in MDA (Figure S1A-D). In addition, for the LL-Salt combined treatment, the changes of these four enzymes were vague. This evidence consistently suggests that the antioxidant system enzymes are an essential factor that causes the differences in H2O2 between two light conditions.

We first performed transcriptome analysis to illustrate further the mechanism of physiological changes in sheepgrasss leaves induced by LL-Salt treatment. Results suggest that the number of clean reads across 18 samples is 65,764,000, accounting for 99.3% (Table S2). The Q30 values across 18 samples were 0.97 (Table S2). Due to limited genome information for sheepgrass, we performed no-references comparisons by mapping the reads to nine genome databases to increase the accuracy of the mapping approach. Results show most reads were fine-mapped to the sequence from the Nr database with 21.6% mapping rates, followed by Pfam, Swiss-prot, and gene-ontology blast (Figure S2A). Similarities analysis suggests that samples within the group for each treatment show a high correlation coefficient with *r*>0.87. Consistent with this, principal component analysis indicates that samples in each group were clustered together except for one sample in the ML_50 group (Figure 2A). The log_2_(FPKM) values across the 18 samples were all around 2, and the distribution of the FPKM values yielded normal distribution (Figure S2C-D). These results confirmed the good quality of the samples used for transcriptome analysis.

**Figure 2.**
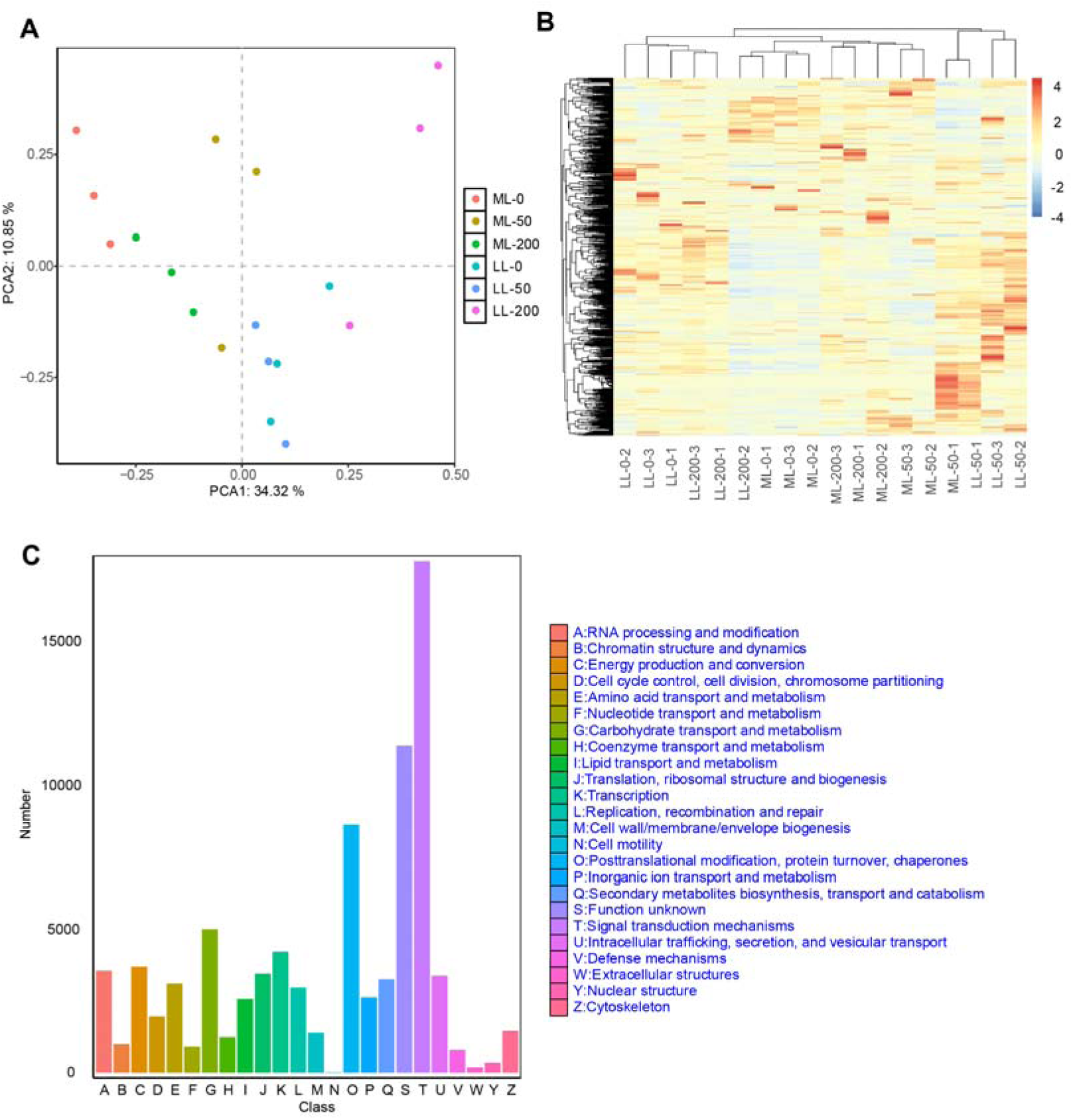
Transcriptome analysis on the grass exposed to either LL or high salt treatment. **A**, Principal component analysis on the global gene expression of sheepgrass leaves exposed to different light and salt treatments. **B**, Heatmap representing the relative abundance of the global gene in 18 samples. **C**, Annotated information of biological pathways of the global gene.

In this regard, we identified 43,562 genes across the 18 samples (visualized in Figure 2B), and most of the genes were annotated to signal transduction mechanism, post-translational modification, protein turnover, chaperones except for functional unknown due to limited gene information (Figure 2C). In addition, very few genes were annotated to extracellular and nuclear structure (Figure 2C). We then compared the LL-induced changes in global gene expression between the ML_0 vs. LL_0 group. There are 4921 downregulated and 4552 upregulated genes with significant differences (DEGs), visualized in Figure 3B. The top 1% of DEGs are listed in Table S3. GO and KEGG analysis on the 4921 downregulated DEGs suggest that the cellular function terms of plastid, plastid part, membrane part, and thylakoid part were significantly enriched in GO analysis. The pathways of starch and sucrose metabolism, photosynthesis, glyoxylate, dicarboxylate metabolism, and carbon metabolism were significantly enriched in the list of KEGG analyses (Figure 3D). Regarding upregulated DEGs, we found that protein-containing complex, non-membrane-bounded organelle, and intracellular organelle lumen were enriched considerably based on GO analysis (Figure S3A). In contrast, the pathways of the ribosome, valine, leucine, and isoleucine biosynthesis, peroxisome were significantly enriched based on KEGG analysis (Figure S3B).

**Figure 3.**
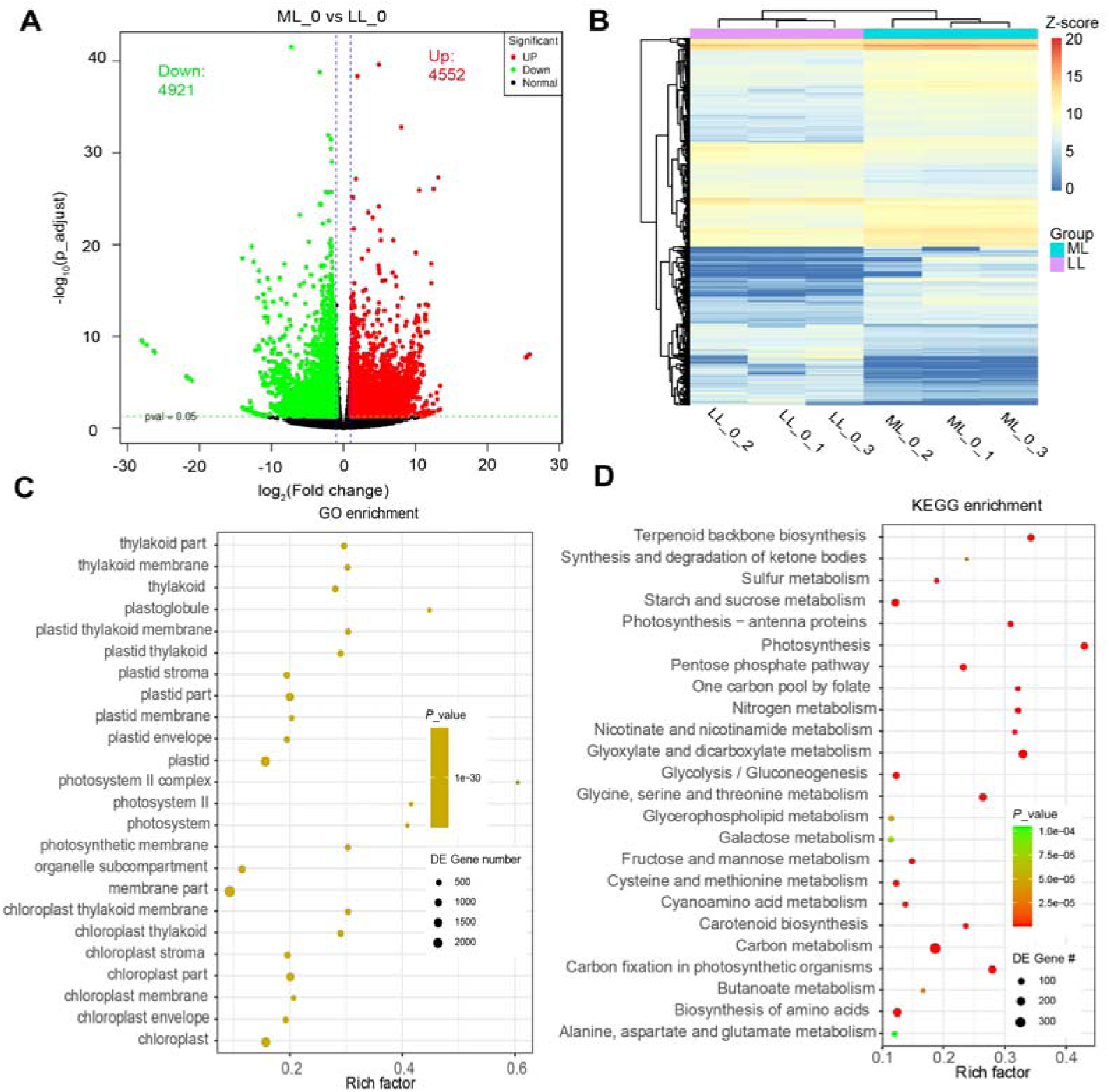
Low light-induced global expression changes in the grass. **A,** Volcano plot representing the differentially expressed genes in comparison of ML_0 and LL_0. **B**, heatmap showing the relative abundances of transcripts in six sheepgrass samples exposed to either LL or ML. **C-D,** GO (**C**), and KEGG (**D**) analysis on the list of downregulated DEGs in LL_0 compared it to ML_0.

Next, we analyzed the DEGs induced by two salt conditions (50 and 200 mM NaCl). Results suggest that 2210 DEGs were overlapped between the comparisons of ML_200 vs ML_0 and ML_40 vs ML_0 (Figure 4A). In the overlapped 2210 DEGs, we found 924 upregulated and 1286 downregulated DEGs in comparison of ML_200 vs ML_0, while 875 upregulated and 1335 downregulated DEGs in comparison of ML_50 vs ML_0 (Figure 4B). Furthermore, we identified 301 common upregulated DEGs and 194 common downregulated DEGs in salt-induced comparisons (ML_200 vs ML0 and ML_50 vs ML_0) for the following GO and KEGG analysis (Table S4). GO and KEGG analysis on downregulated DEGs suggest that the pathways of protein phosphorylation, photosynthesis, and phosphate metabolic process were significantly enriched based on GO analysis.

**Figure 4.**
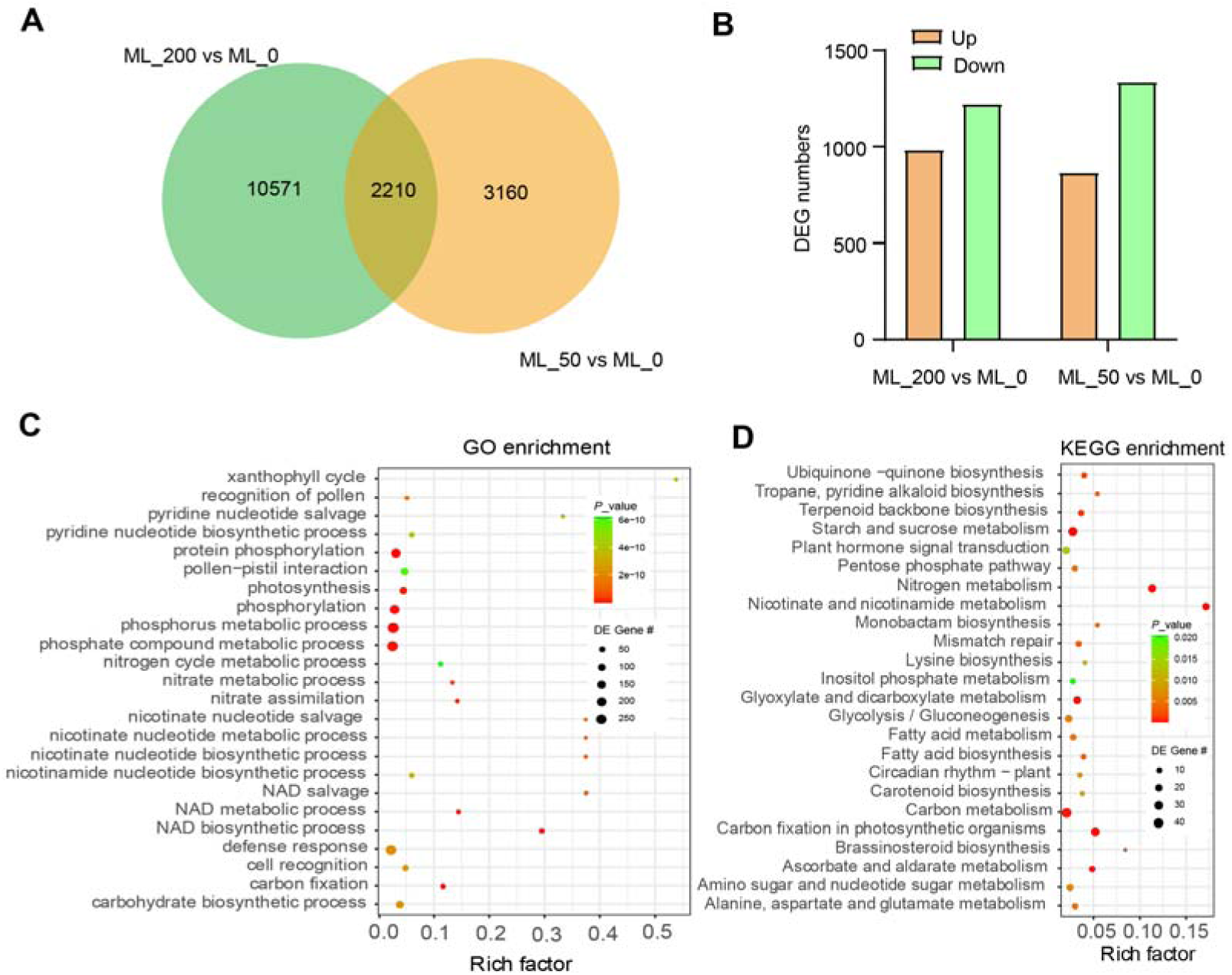
Salt-induced global gene expression changes in sheepgrass. **A**, Venn diagram showing the common differentially expressed genes in the comparisons of two salt treatments (ML_200 vs ML_0 and ML_50 vs ML_0). **B**, Summary of differentially expressed genes with either upregulated or downregulated patterns in comparing two salt treatments. **C-D**, GO (**C**), and KEGG (**D**) analysis on the list of downregulated DEGs in both salt treatments (ML_200 and ML_50) relative to ML_0.

In contrast, starch and sucrose metabolism, pentose phosphate pathways, glyoxylate and dicarboxylate metabolism, and carbon metabolism were enriched considerably based on KEGG analysis (Figure 4C-D). Regarding upregulated DEGs induced by salt treatment, we found that the pathways of response to stimulus, response to stress, response to external stimulus, and defense response were significantly enriched based on GO analysis. In contrast, based on KEGG analysis, starch and sucrose, glutathione metabolism, arginine and proline interaction, and galactose metabolism were significantly enriched in the upregulated DEGs (Figures S4A-B).

We then analyzed the DEGs induced by salt and light interaction effects. Results suggest that 16 overlapped DEGs were identified in both light (including ML_0 vs. LL_0) and salt (both ML_200 vs. ML_0 and ML_50 vs. ML_0) groups (Figure 5A). We named these as LL-Salt responsive genes listed with their relative transcript abundance in Figure 5B. Furthermore, the six genes with different expression patterns in comparisons of ML_200 vs ML_0, ML_50 vs ML_0, and LL_0 vs ML_0 were selected to conduct qPCR. The results confirm that expression levels of a phytochrome-interacting transcription factor 4 (PIF4) and a WD repeat-containing protein 44 (*WDRCP44*) were increased by LL. In contrast, receptor kinase-like protein Xa21 (*Xa21*), peroxidase 5 (*POD5*), and receptor protein kinase (*ZmPK1*_like) were decreased with no significant changes in ABC-transporter genes (Figure 5B-G). Regarding salt stress, results confirmed that the expression level of *PIF4*, *Xa21*, *POD5*, *ABC-transporter,* and *ZmPK1-like* was decreased, while *WDRCP44* was increased by salt stress (Figure 5C-G).

**Figure 5.**
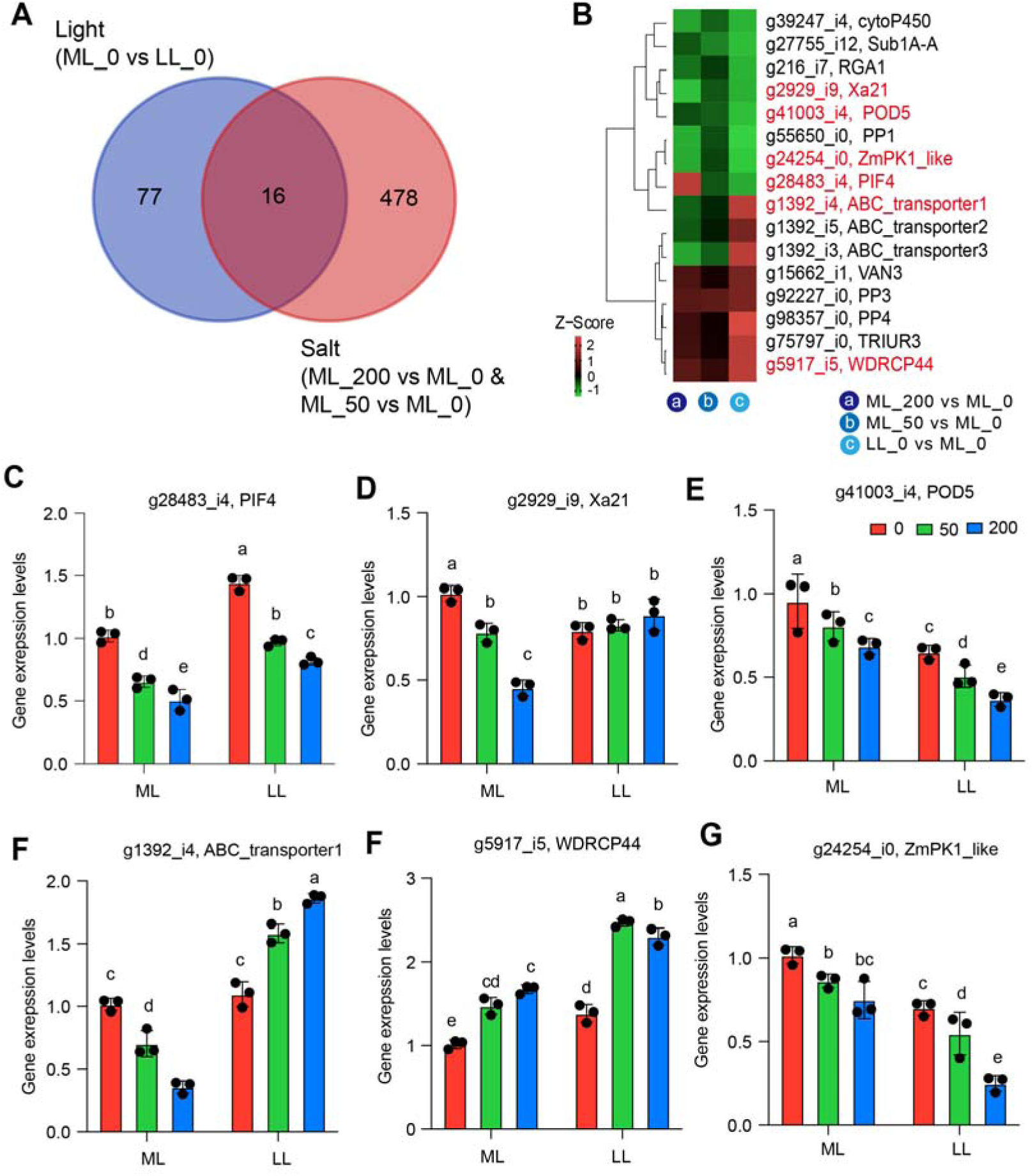
Interactive effects of salt and light on global gene expression in sheepgrass based on transcriptome analysis. **A**, Venn diagram showing the overlapped genes in comparisons of light (ML_0 vs LL_0) and two salt treatments (ML_200 vs ML_0 and ML_50 vs ML_0). **B**, The relative abundance of the overlapped 18 genes. **C-G**, Relative expression levels of genes involved in the overlapped gene list in sheepgrass leaves exposed to light or salt treatment. Each bar data represents the mean of replicates (*n*=3). Different letters represent significant differences based on one-way *ANOVA* followed by Tukey’s HSD tests (*P*<0.05).

Furthermore, we performed a non-targeted metabolism analysis to identify the DAMs in the six combinations of light and salt stress in sheepgrass. Similarities analysis suggests that samples within the group for each treatment show a high correlation coefficient with *r*>0.92. Principal component analysis (PCA) indicates consistently that samples in each group were clustered together (Figure 6A). Most metabolites were identified to annotate in fatty acids, organ-oxygen compounds, carboxylic acids, and derivatives (Figure S5B). Volcano plots show that 65 upregulated DAMs and 56 downregulated DAMs in comparison of ML_200 vs ML_0 (Figure 6B; Table S6).

**Figure 6.**
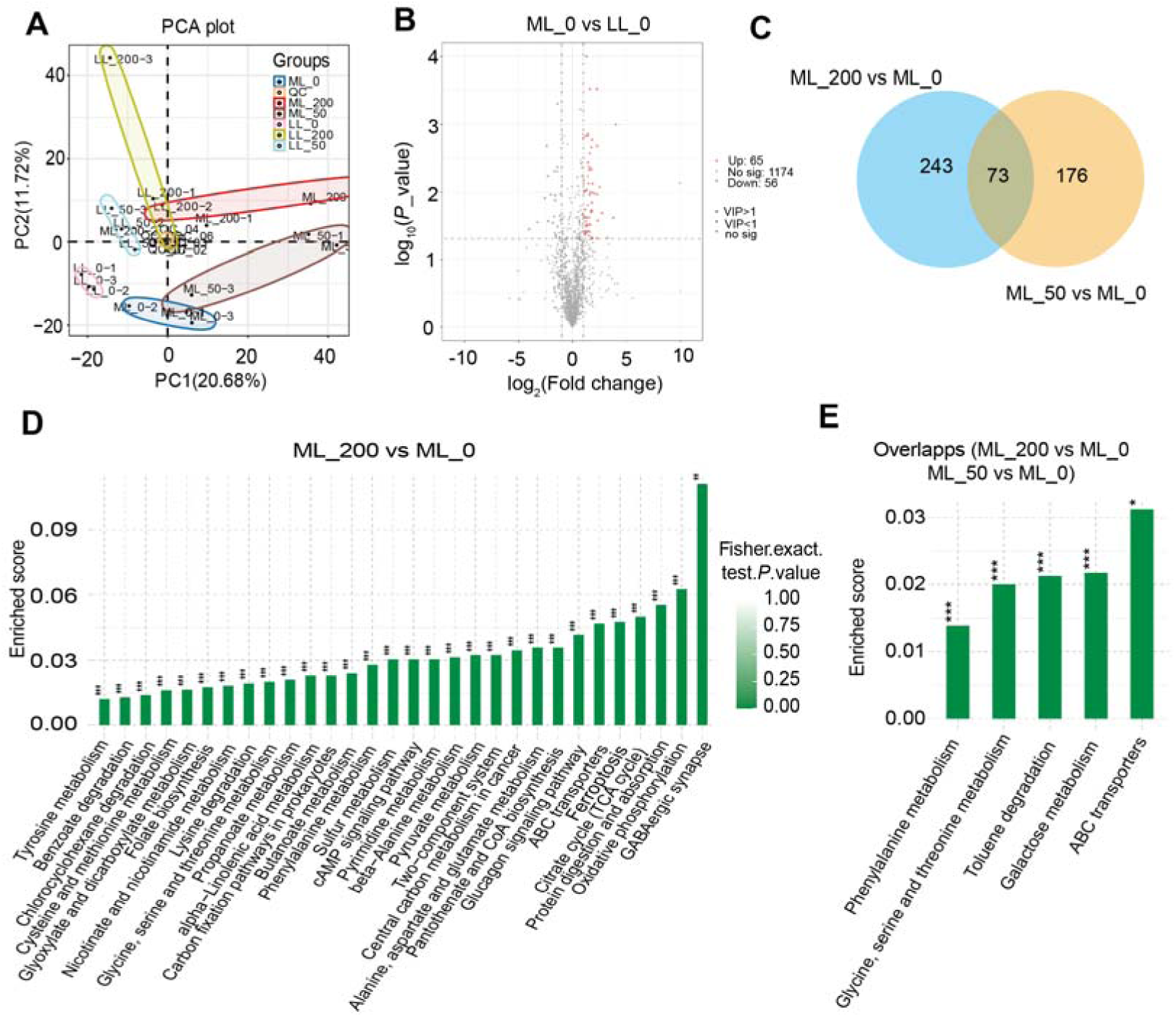
Differentially abundant metabolites in sheepgrass exposed to either salt or light conditions. **A**, Principal component analysis on the 1254 identified metabolites across 18 sheepgrass leave samples. **B**, Volcano plot representing the differentially abundant metabolites in comparison of ML_0 vs LL_0. **C**, Venn diagram showing the overlapped differentially abundant metabolites in two salt treatments (ML_200 vs ML_0 and ML_50 vs ML_0). **D-E,** KEGG analysis on the list of differentially abundant metabolites in comparison of ML_200 vs ML_0, in comparison of two salt treatments (ML_200 vs ML_0 and ML_50 vs ML_0).

Salt stress-induced DAMs were then analyzed. Venn diagram shows that 73 overlapped DAMs between the comparisons of ML_200 vs ML_0 and ML_50 vs ML_0 (Figure 6C; Table S7). KEGG analysis on the list of 278 DAMs in comparison of ML_200 vs. ML_0 suggest that metabolic pathways of GABAergic synapse, protein digestion and absorption, citrate cycle (TCA) cycle, ABC transporter, pyruvate metabolism, glyoxylate, and dicarboxylate metabolism were significantly enriched (Figure 6D). Furthermore, KEGG analysis on the 35 overlapped DAMs shows phenylalanine metabolism, glycine, serine and threonine metabolism, galactose metabolism, and ABC transporter (Figure 6E).

In addition, we identified 16 common DAMs between the comparisons of light response (ML_0 vs LL_0) and salt response (ML_200 vs ML_0 & ML_50 vs ML_0) (Figure 7A-B; Table S8). Among them, three identified metabolites involved in the TCA cycle were all increased in LL treatment compared to ML, while downregulated in two salt stress treatments (ML_200 and ML_50) relative to ML_0 (Figure 7B; Table S8). For the carbon metabolism pathway, we identified that 7 DAMs, including glucose, raffinose, sucrose, D-ribose, D-fructose, D-mannose, and ribulose 5-phosphate, were all downregulated in both light treatment (ML_0 vs LL_0) and salt treatments (ML_200 vs ML_0 & ML_50 vs ML_0). In addition, an identified metabolite, glutathione disulfide, in the GSH/GSSG redox system together with two photorespiratory pathway intermediates, i.e., serine and glycolate, were all downregulated in LL treatment, while upregulated in the two salt treatments (Figure 7B; Table S8).

**Figure 7.**
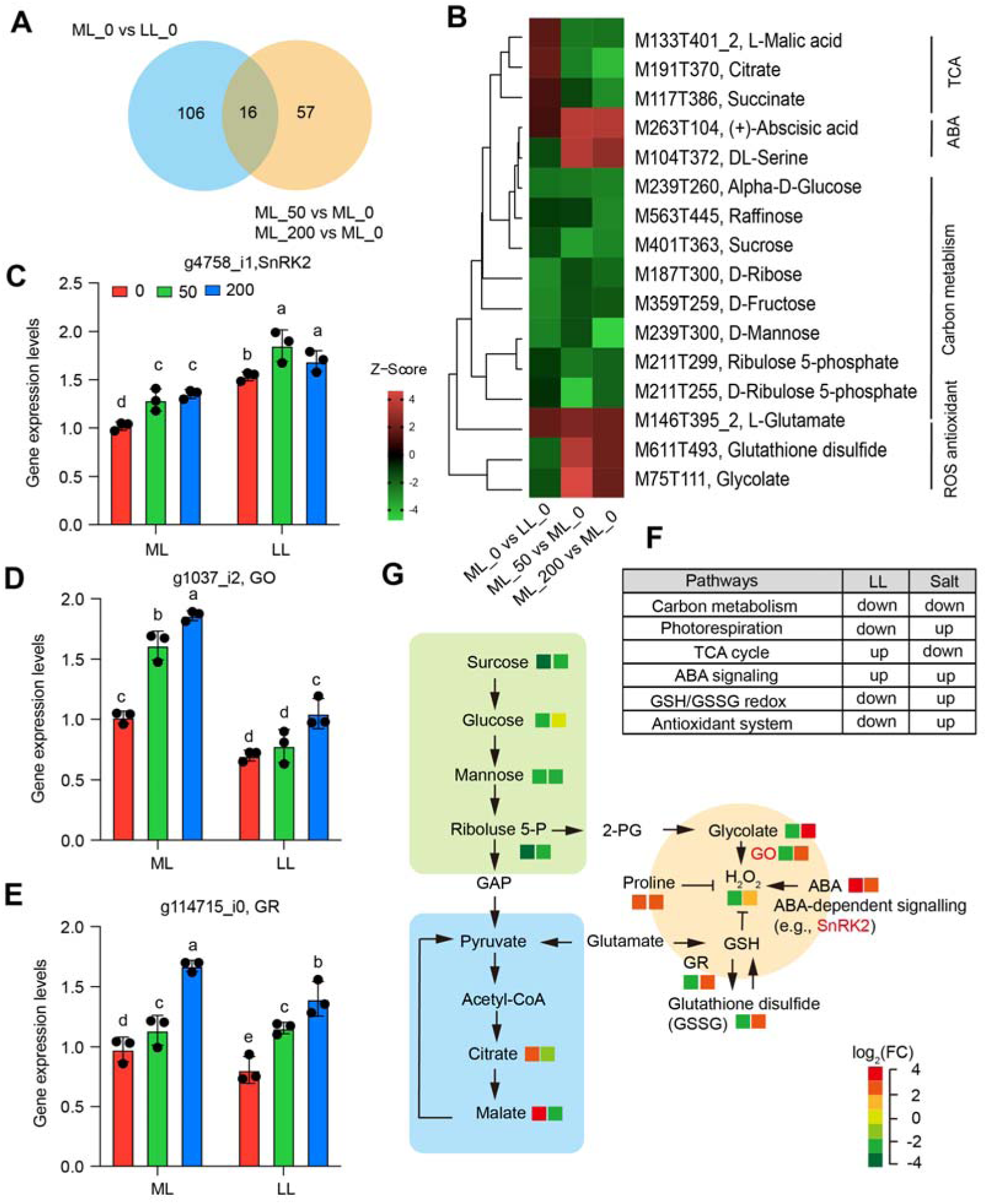
Summary of the key metabolic pathways that participate in the response of low light and salt stress in sheepgrass. **A,** Venn diagram of differentially abundant metabolites in comparisons of salt and light treatments. **B,** Heatmap representing the relative abundance of overlapped differentially abundant metabolites in comparisons of light (ML_0 vs LL_0) and salt (ML_200 vs ML_0 & ML_50 vs ML_0). **C-E,** Relative expression levels of genes involved in ABA signaling pathway, photorespiration, and GSH/GSSG redox system pathways in sheepgrass leaves exposed to either LL or salt treatments. **F,** Summary of regulation pattern of key metabolic pathways in response to low light and salt stress in sheepgrass leaves. **G**, Working model showing the interactive effects of LL and salt on key metabolic pathways in sheepgrass leaves.

We measured the expression levels of some regulatory genes to validate the patterns of metabolites involved in these metabolic pathways. Results show that the expression levels of serine/threonine-protein phosphatase 2 (*SnRK2*) in the ABA signaling pathway were significantly increased in both LL treatment and two salt treatments (Figure 7C). The glycolate oxidase gene (*GO*) expression levels in the photorespiration pathway were decreased considerably by LL treatments but significantly increased in two salt treatments (Figure 7D). In the GSH/GSSG system, we found that glutathione reductase (*GR*) expression levels were significantly decreased by LL treatment but increased considerably by two salt treatments (Figure 7E). The expression levels of three genes were in line with the metabolite’s patterns (Figure 7C-E).

In summary, carbon metabolism pathways were downregulated in both LL treatment and salt treatments. Photorespiration was downregulated and upregulated. TCA cycle was upregulated and downregulated in LL and salt treatment, respectively. ABA signaling was upregulated for both LL and salt treatments, and the GSH/GSSG redox system was down and upregulated in LL and salt treatments, respectively (Figure 7F). Some intermediates involved in the antioxidant system, including H_2_O_2_, decreased catalase activities under LL while increasing in salt treatments (Figure S1A; Figure 7F). Hence, we proposed the interactive pattern of primary metabolic pathways in response to LL and salt (Figure 7G). Sucrose contents were depredated for either LL or salt treatment, while LL speeds up, and salt inhibits the TCA cycle (Figure 7G). Salt stimulates photorespiration, leading to the accumulation of H_2_O_2_, together with ABA signaling, proline, and GSSG accumulation (Figure 7G). In contrast, LL inhibits photorespiration, hence resulting in less production in H_2_O_2_, collectively working with GSSG reduction (Figure 7G).

## Discussions

LL always occurs in natural conditions, which affects the growth of plants from molecular to morphological[7–9]. Extreme LL, especially with other stressors such as salinity, is the most challenging for plants that respond dynamically to these conditions [10]. Sheepgrass is a promising model material for salt tolerance study [2]. However, the mechanism underlying such a LL-Salt combined process remains unclear. Meeting the increasing demand for higher productivity and high protein content of sheepgrass, new ways will be required for sheepgrass molecular breeding in the future. Biomass under LL-Salt determines carbohydrate gains and defense responses [21]. Thus, there is a pressing need to understand the molecular mechanism of biomass development under LL-Salt. In this study, we demonstrated that several metabolic pathways are essential for adaption to LL-Salt in sheepgrass. Both LL and salt treatment induce downregulation of carbon metabolism and upregulation of the ABA signaling pathway. The two stresses have their separate regulatory pathways, as well, and they especially function differently in cellular redox homeostasis.

### Downregulation of Carbon metabolism in LL-Salt combined condition

In plants, sugars serve as metabolic resources and structural components of cells, and they undergo osmotic adjustment under various stress conditions, including salt stress [22–25]. Sucrose is also the main source of carbon and energy for plant metabolism. Plants initiate regulatory mechanisms under abiotic stress conditions by adjusting sucrose content and promoting carbohydrate redistribution to adapt to such stress conditions [26]. Consistent with the observation in our study, we found that reduction in sucrose as well as other metabolites in the carbon metabolic pathway under combined stress condition of LL-Salt, including fructose, mannose, and ribose, suggesting that sucrose in the leaves tended to degrade sugar and glucose [26], leading to the hydrolysis of sucrose into small molecular carbohydrates for adaption to the both LL and salt stress. Similar to those found in salt stress conditions, in LL, the reduction in the sucrose content also exacerbates the decline in photosynthetic carbon assimilation [27] indicated by the reduction in photosynthetic efficiency, including Fv/Fm, which inhibit energy-containing substances such as ATP [28], with photorespiration reduction [29].

### Photorespiration downregulated by LL but upregulated by salt stress

As a bypass pathway of carbon metabolism, photorespiration, an inevitable process in photosynthesis, plays a supporting role not only in photosynthetic CO_2_ assimilation [30] but also in ROS production. Importantly, the growth LL intensity significantly affects the development of the photorespiratory pathway [31]. Although photorespiratory intermediates such as glycolate and glycerate inhibit the Calvin cycle [30], they can be converted to glycerate-3-phosphate through the photorespiratory pathway [32]. This process is critical for photosynthesis and photoprotection [33], suggesting its potential dual function in mediating carbon assimilation metabolism. Very interestingly, salt stress could accelerate photorespiration, which is totally different from what happens under LL (Figure 8F-G), which is possibly attributed to its important function for preventing free radicals (ROS) accumulation from maintaining cellular redox homeostasis [34].

### Damage repair and defense response under salt stress

Oxidative stress caused by excessive ROS is a direct indicator reflecting cellular damage caused by salt stress [35]. In order to determine whether antioxidant enzymes can promote ROS scavenging and mitigate the oxidative damage under LL-Salt combined stress in sheepgrass leaves, we measured the activities of SOD, POD, MDA and CAT. In general, POD, SOD, MDA, and CAT are plant-specific enzymes involved in lignin formation, the cross-linking of cell wall components, and the removal of H_2_O_2_ against abiotic stresses [36, 37]. Our results showed that the activities of CAT and POD were decreased due to salt stress but showed an opposite pattern in response to LL (Figure S1A-B), while activities of MDA and SOD were increased by salt, and two enzymes showed similar or increase in LL response. This evidence suggests that different antioxidants have different roles in response to salt and LL, although a common function for the removal of H_2_O_2_ is needed when plants experience LL-salt combined stress in sheepgrass leaves. This might be associated with an adaptive response in plants to such conditions, as indicated previously [38, 39]. Meanwhile, many studies have shown that both LL and salt stress-induced ROS accumulation might be a mechanism to protect plants rather than cause damage, at least at the initial stage [37, 40, 41].

### Roles of GSG/GSSH redox system in removal of ROS under LL-Salt

In the antioxidant scavenging system, the ascorbate-glutathione cycle represents a central component pair of the redox system participating in adjusting growth and development to changing environmental conditions, including alterations in the intensity and spectrum of light [42–44]. Notably, glutathione reductases (GR) mediate the conversion of GSSG into GSH and play a crucial role in the elimination of H_2_O_2_ molecules through the ascorbate–glutathione cycle. With decreasing light intensity, the amount of its reduced (GSH) and oxidized (GSSG) form and the GSSG/GSH ratio decreased. It was reported that the shaded leaves of crops, like soybean, had a much lower glutathione reductase (GR) activity and reduced glutathione (GSH) level compared to those grown in full light [45, 46]. Consistent with this, our results suggest that the expression levels of glutathione reductase in sheepgrass leaves were also significantly decreased in LL treatments, leading to higher GSSG levels than that under ML.

### Effects of LL-salt combined stress on TCA cycle

The citrate cycle is one of the most important metabolic pathways, and it provides major energy resources, such as ATP and NADPH, for growth development [47]. In the initial step, pyruvate is converted to acetyl-CoA from mitochondria to chloroplast and initiates TCA with produce many amino acids, including citrate, succinate, and malate. In particular, Malate and citrate underpin the characteristic flexibility of central plant metabolism by linking mitochondrial respiratory metabolism [48]. Our metabolism analyses showed that all three metabolites were upregulated in LL condition but downregulated in salt stress (Figure 7B). The reduction in the three metabolites under salt stress suggests oxidative stress occurs, as observed in another study in barley [49]. In addition, the TCA cycle functions in the light at a rate similar to that in the dark except for a brief initial inhibition on the transition from dark to light [50], suggesting greater effects of salt stress than that under LL conditions. Taken together, our results suggested that the TCA cycle plays a key role in response to salt stress by regulating the abundance of specific metabolites, as observed previously in other species [51].

## Declaration Abbreviations

Not applicable.

## Ethics approval and consent to participate

Our rice collection work complies with the laws of the People’s Republic of China and has a permission letter from Institute of Grass Research, Heilongjiang Academy of Agricultural Sciences. Voucher specimens were identified by Prof. Wei Li (Heilongjiang Academy of Agricultural Sciences) and kept at Heilongjiang Rice Quality Improvement and Genetic Breeding Engineering Research Center (No: SG001-SG058). All methods were carried out in accordance with relevant guidelines and regulations.

## Consent for publication

Not applicable.

## Acknowledgements

Not applicable.

## Author contributions

MQ and JL planned and designed the research. JL, SF, HZ, ZI, CS, WH, KH, XM, MD, ZL abd BG performed experiments. JL, SF and HZ analysed data. JL, MQ and ZI wrote the manuscript.

## Data and materials availability

All data is available in the manuscript or the supplementary materials.

