## Supplementary material for "Integrative analysis of transcriptome and metabolism reveals functional roles of redox homeostasis in low light and salt combined stress in *Leymus chinensis*": Supplementary Figures.docx

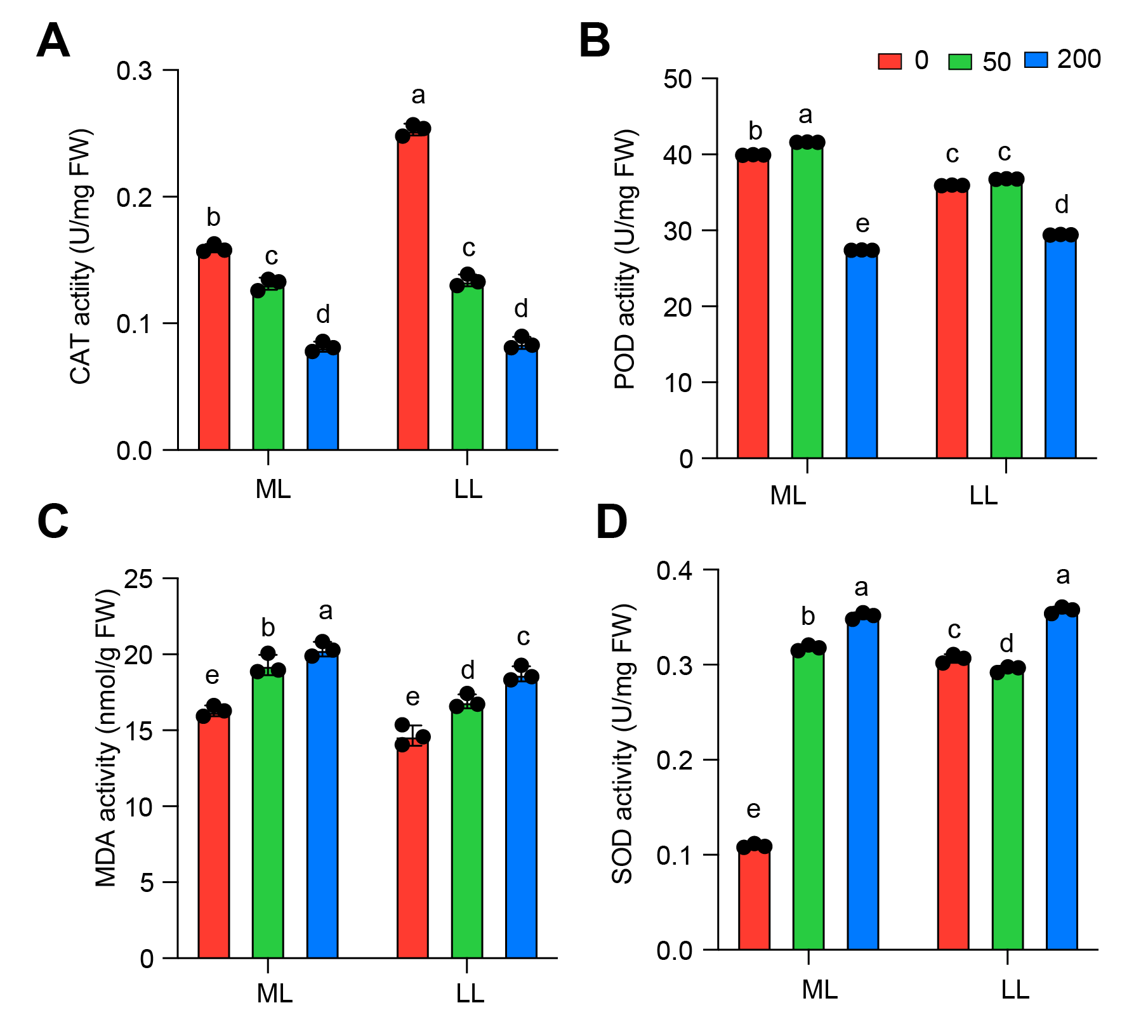

**Figure S1.** Antioxidant enzyme activities in grass exposed to either LL or high NaCl treatment. **A,** Catalase (CAT) activity; **B,** Peroxidase (POD) activity. **C,** Malondialdehyde (MDA) activity; **D,** Superoxide dismutase (SOD) activity. Each bar data represents the mean of replicates (*n*=3). Different letters represent significant differences based on one-way *ANOVA* followed by Tukey’s HSD tests (*P*<0.05).

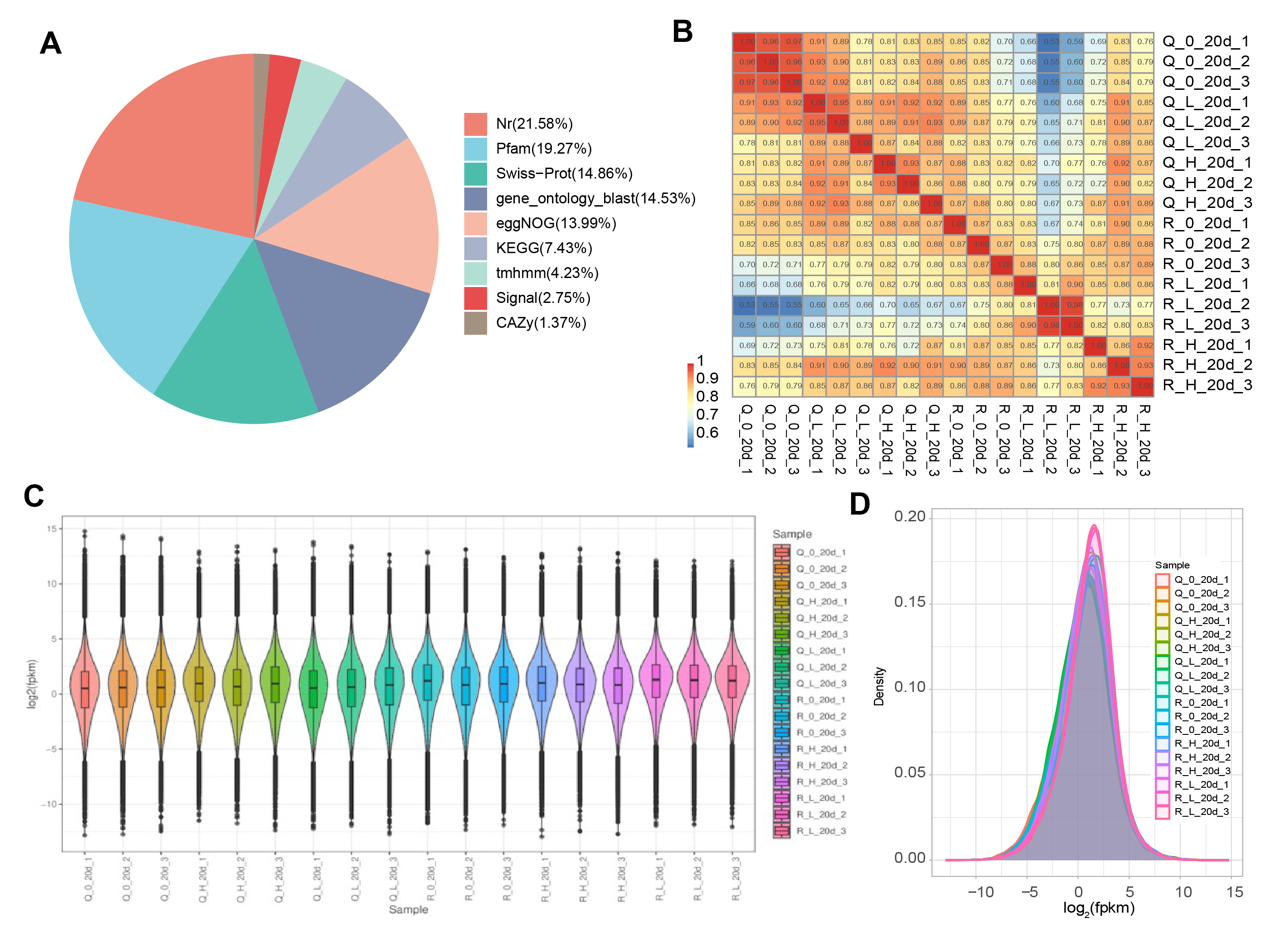

**Figure S2.** Quality control of transcriptome dataset in leaves of sheepgrass exposed to either LL or high NaCl. **A,** Distribution of genome database that mapped by transcripts. **B,** Similarity analysis on the transcript abundances in 18 samples. **C,** Distribution of log_2_(FPKM) values across 18 samples. **D,** histogram of log_2_(FPKM) values across 18 samples.

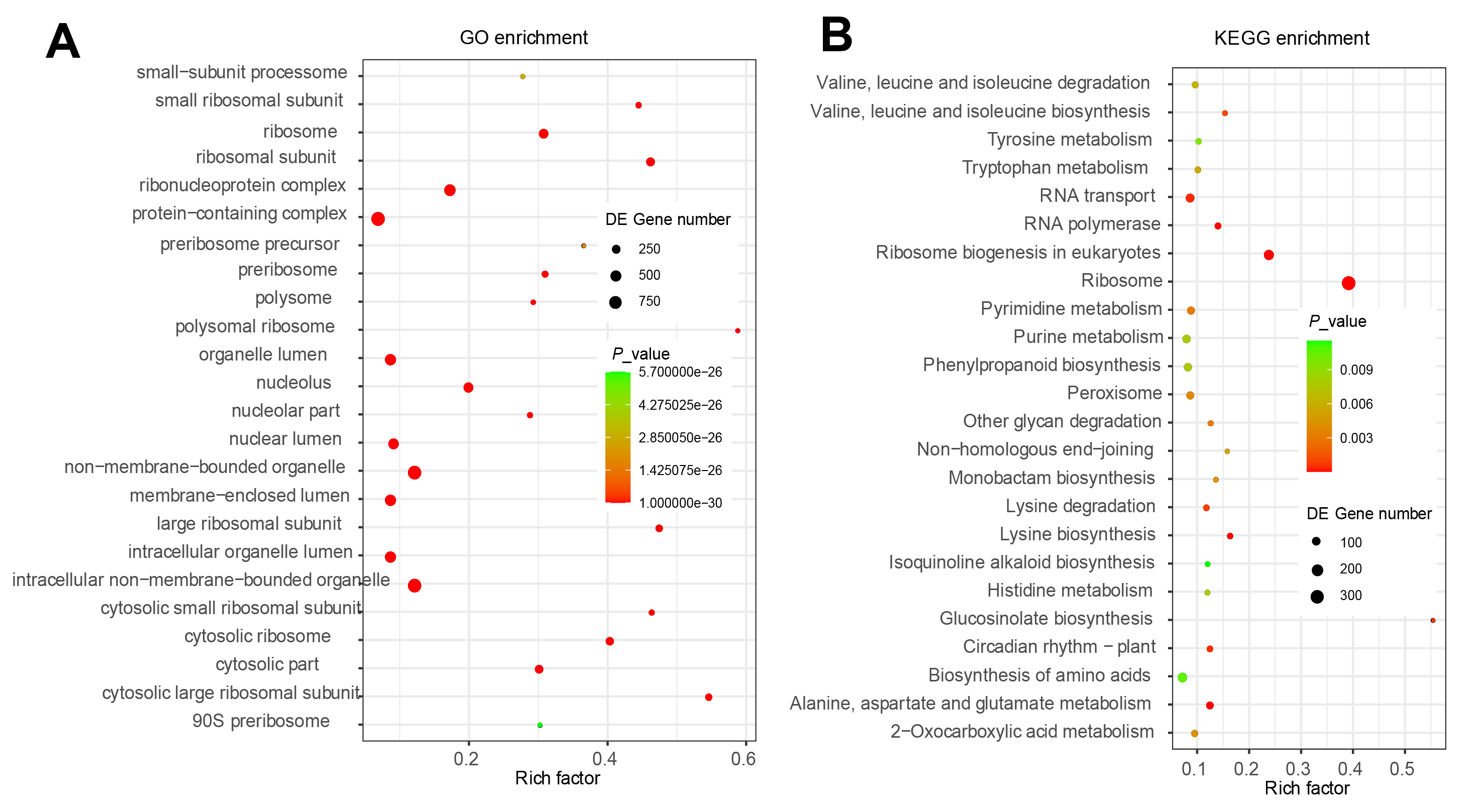

**Figure S3.** GO and KEGG analysis on the list of upregulated DEGs in LL_0 compared it to ML_0. **A**, GO; **B**, KEGG.

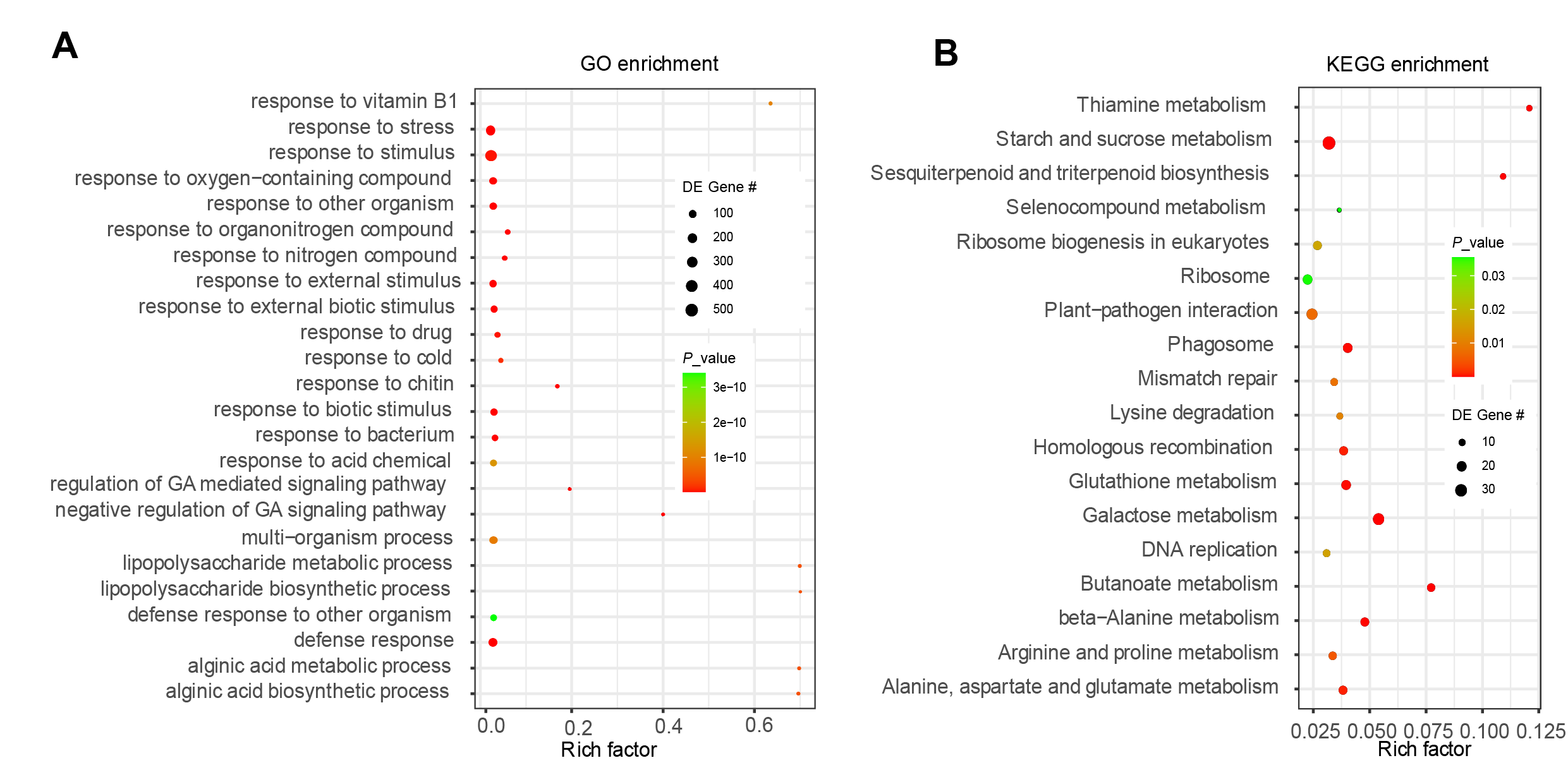

**Figure S4.** GO and KEGG analysis on the list of upregulated differentially expressed genes in two salt treatments (ML_200 vs ML_0 and ML_50 vs ML_0). **A,** GO. **B,** KEGG.

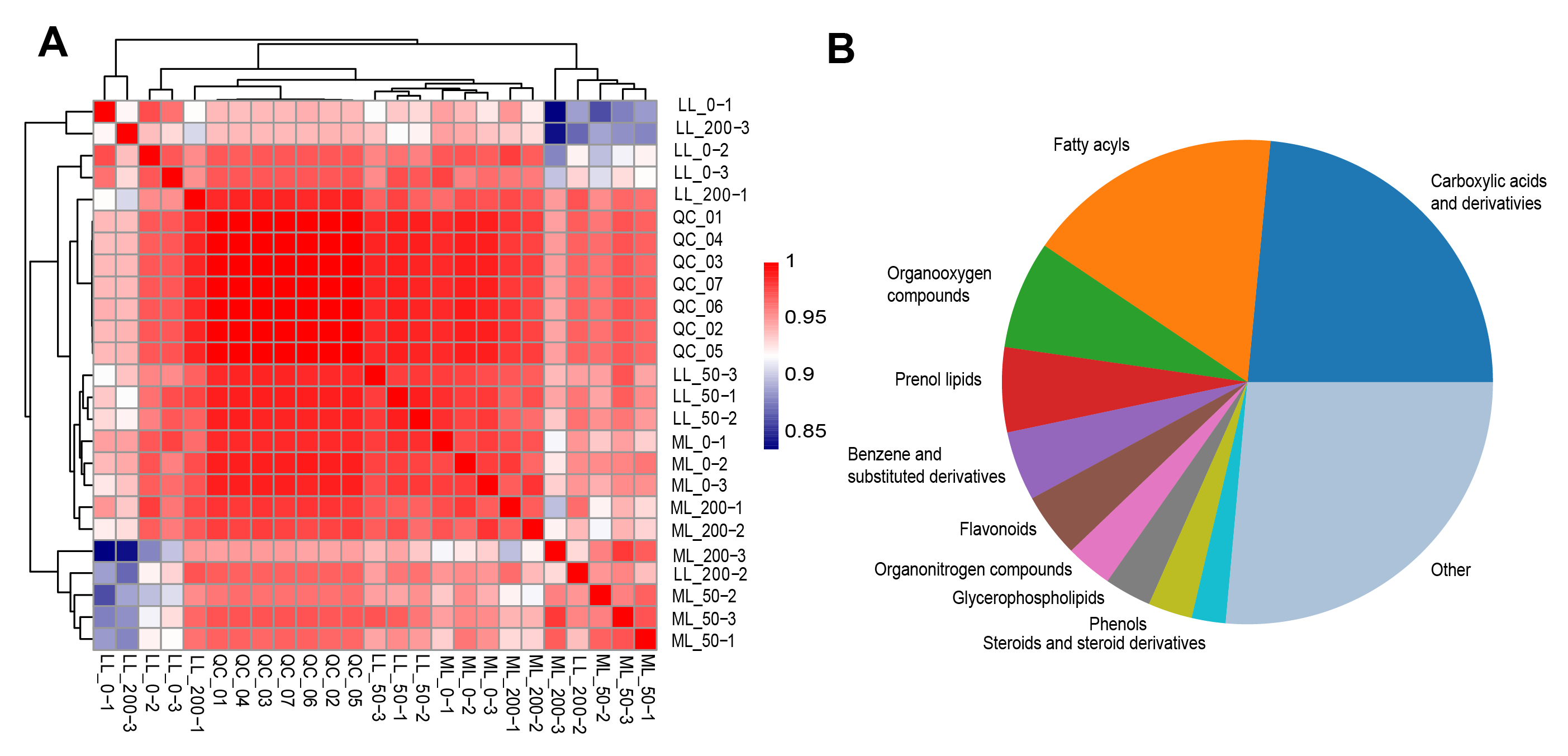

**Figure S5.** Quality control of non-targeted metabolites analysis. **A,** Similarity analysis on the transcript abundances in 18 samples. **B,** Functional analysis of KEGG database that mapped by different metabolites across 18 sheepgrass leaves samples.
